# Using stationary vital rates in impact assessments may underestimate potential threat

**DOI:** 10.1101/2021.07.01.450685

**Authors:** C. Horswill, J.A.O. Miller, M. J. Wood

## Abstract

Population viability analysis (PVA) is commonly used to assess future potential risks to threatened species. These models are typically based on mean vital rates, such as survival and fecundity, with some level of environmental stochasticity. However, the vital rates of wild populations, especially those already exhibiting declining trajectories, may be nonstationary, such that the mean or variance changes over time. In this study, we examined whether including observed temporal trends in vital rates affects the predictive accuracy of PVA, as well as the projected impact associated with a hypothetical threat. To achieve this, we ran a series of simulations using Leslie matrix PVA models that included different combinations of environmental stochasticity, temporal trends in vital rates, and threat. We apply our analysis to a long-lived colonial species of seabird, the black-legged kittiwake *Rissa tridactyla*, that is classified as globally Vulnerable and is potentially highly sensitive to offshore renewable energy development. We found that including observed temporal trends in vital rates was (i) crucial for the accurate reconstruction of observed population dynamics and (ii) had a dramatic effect on the projected impact from the hypothetical threat. In an era when many animal and plant populations are declining due to long-term trends in their vital rates, we identify that including this demographic structure is essential for robustly evaluating potential threats using PVA models. Omitting observed temporal trends in vital rates from impact assessments is highly likely to yield unreliable results that could misinform conservation and management decision making. This result has immediate application for conducting impact assessments on protected species and populations.

## Introduction

In many countries, it is necessary to conduct a full evaluation of potential impacts to protected wildlife populations to gain consent for certain activities, such as biological resource use, land-use change or energy production (e.g., EU Birds Directive 2009/147/EC, EU Habitat Directive 92/43/EEC, EU EIA Directive 85/337/EEC). A commonly used approach for inferring how populations may respond to these activities is to predict how vital rates – such as survival or fecundity – are likely to be impacted, and then use these altered values to evaluate the population response. Population viability analysis (PVA) provides an analytical framework for conducting such impact assessments (Reed et al. 2002).

The analytical structure of a PVA does not always include a demographic model (Reed et al. 2002), however applications used in impact assessment and conservation decision making typically require a quantitative approach (Beissinger & Westphal 1998; Chaudhary & Oli 2020; Searle et al. 2020). Accurate model structure and demographic data, as well as environmental and demographic stochasticity are widely acknowledged as important factors for improving the predictive accuracy of PVA (Beissinger & Westphal 1998; Coulson et al. 2001; Reed et al. 2002; Pe’er et al. 2013; Chaudhary & Oli 2020). However, temporal trends in vital rates occurring independently of the future threat are often overlooked (Chaudhary & Oli 2020; Searle et al. 2020). This demographic structure is important because it may influence population growth. Furthermore, many populations are experiencing temporal trends in their vital rates (e.g., Fig. 1) in response to climate change and habitat degradation (e.g. Frederiksen et al. 2004; Sydeman et al. 2021).

**Fig. 1.**
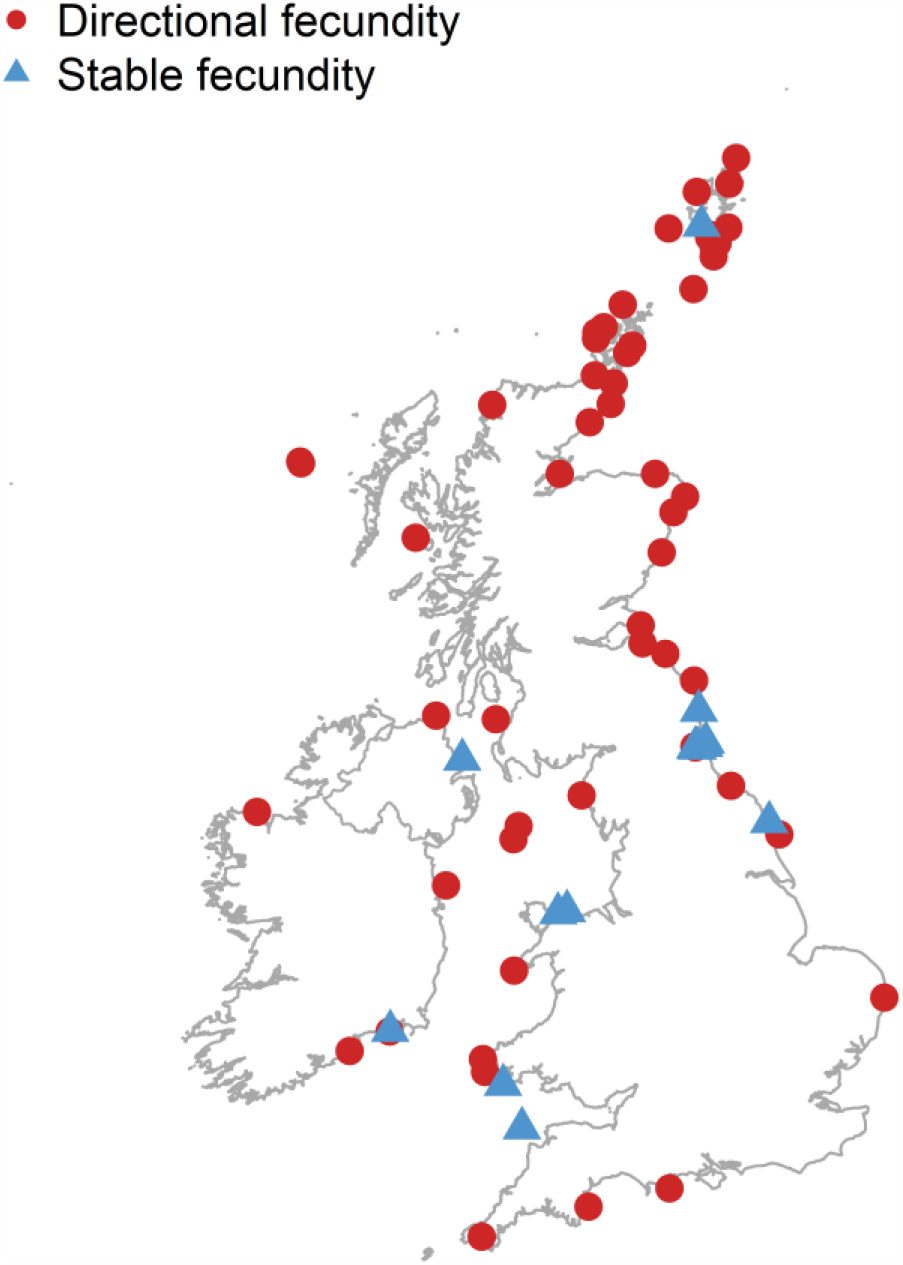
The majority (79%) of black-legged kittiwake colonies in the UK demonstrate a directional temporal trend in their rates of fecundity. Colonies with directional fecundity shown as red circles, those with stable fecundity shown as blue triangles.

In this study, we run a series of Leslie matrix PVA models to examine how observed temporal trends in vital rates influence predictive accuracy, as well as the projected impact associated with a hypothetical threat. We apply our analysis to a species of seabird, black-legged kittiwakes *Rissa tridactyla* (hereafter kittiwakes), that is classified as globally Vulnerable (Birdlife International 2019) and is potentially highly sensitive to additional mortality impacts from collision with offshore wind energy developments (Furness et al. 2013; Bradbury et al. 2014). Evaluating potential impacts to this species is commonly part of the consenting process for offshore wind development in the UK (Ruffino et al. 2020), but currently these assessments do not include temporal trends in vital rates (e.g., Searle et al. 2019). We show that including this structure in impact assessments is essential for obtaining meaningful estimates of potential threats.

## Methods

### Prevalence of temporal trends in vital rates

To demonstrate the potential prevalence of temporal trends in the vital rates of seabirds, we collated colony-specific fecundity data for kittiwakes breeding in the UK. Here, fecundity rates are defined as the proportion of apparent nesting attempts that produce a successful fledgling each year. Data were extracted from the Seabird Monitoring Programme Database (SMP 2020), and we included colonies with at least eight years of data between 1985 and 2019. To identify colonies with a significant temporal trend in rates of fecundity, we used Poisson generalised linear models fitted to each colony with a log link function. In these models, the response variable was set as the number of chicks alive at fledging for each colony, and the number of nests was included on the log scale as an offset. This analysis was conducted in Program R (v. 4.0.2) (R Core Team 2020). To examine whether kittiwakes demonstrate a temporal trend in rates of fecundity at the national level, we fitted a Poisson generalised linear mixed model to the complete dataset using the R statistical package “lme4” (Bates et al. 2016).

### How do temporal trends in vital rates influence predicted population dynamics?

We ran five PVA simulations to examine how observed temporal trends in vital rates may influence the predictive accuracy of PVA, as well as the projected impact associated with a hypothetical threat (Table 1, code provided as supplementary information). The first PVA simulation was deterministic, based on mean demographic rates only. The second model incorporated environmental stochasticity by assigning demographic rates from a beta distribution for survival and a gamma distribution for fecundity. To set the expectation and variance of these distributions, we used colony specific values for kittiwakes breeding on Skomer Island, Wales (51.74°N, 5.30°W). The vital rates for this colony, estimated for the period 1989 to 2018, are 0.86 (±0.06 SD) for adult survival (*ϕ*_*t*_) and 0.68 (±0.08 SD) for fecundity (*f*_*t*_) (Horswill et al. *in review*). The third model was identical to the second model but applied an additional 5% annual mortality to each age class to simulate the hypothetical threat. The fourth model also included environmental stochasticity but added a negative temporal trend on fecundity corresponding to that observed at Skomer (slope = -0.02, 95% credible interval: -0.04, 0.00, Horswill et al. *in review*). Finally, the fifth model included environmental stochasticity, the negative temporal trend on fecundity, and the hypothetical threat, i.e., an additional 5% annual mortality applied to each age class (Table 1).

**Table 1.**
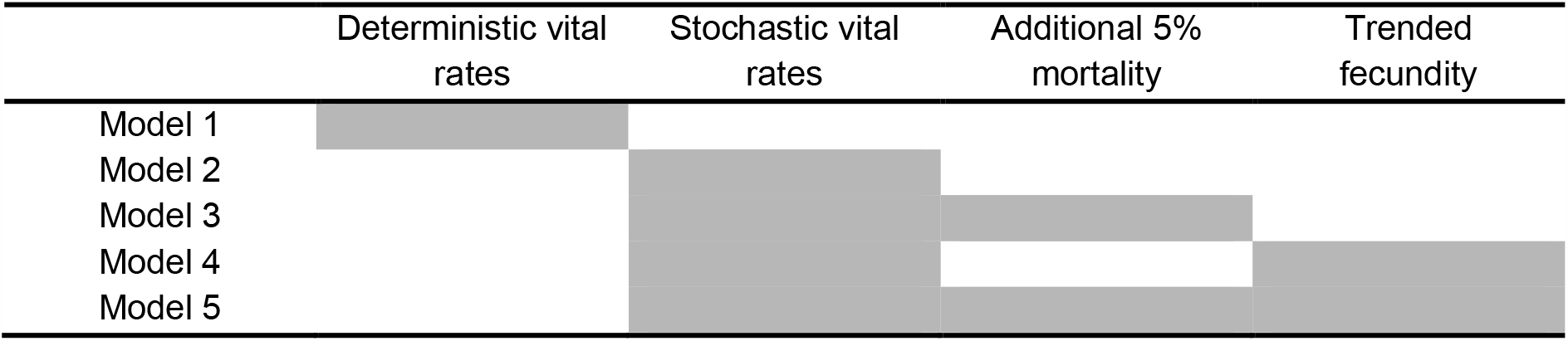
Experimental design for the five PVA simulation models.

For each PVA simulation, we used a Leslie matrix approach following a pre-breeding census to predict the colony size (*N*_*t*_) at time *t* +1 as a function of colony size at time *t* (Eqn. 1). Previous studies have shown that kittiwakes first breed between three and five years old (Porter & Coulson 1987; Cam et al. 2002), and we included this structure by assuming that 25% of 3 year olds and all birds 4 years or older are breeders (Eqn. 4). This structure has been identified as the most suitable structure for describing the observed population dynamics of kittiwakes breeding on Skomer Island (Horswill et al. *in review*). Like many seabirds, kittiwakes are largely unobservable during the first years of life. Therefore, colony-specific estimates of juvenile (i.e., during the first year post fledging *S*_*t*_) and immature (i.e., before recruitment *I*_*t*_) survival rates are limited (Horswill & Robinson 2015). Available estimates for black-legged kittiwakes breeding in France report fledgling survival rates to be 75% of adults, immature survival rates (i.e., from age 1 to 2 years) to be 87.5% of adults, and survival rates to be comparable to adults from age 2 onwards (Cam et al. 2005). Therefore, we assumed an additive relationship between age classes (Cam et al. 2005; Horswill et al. 2014, 2016), and used the respective scalars to estimate juvenile and immature survival rates from rates of adult survival. We assumed a 1:1 sex ratio in the colony and halved fecundity to model female numbers only.

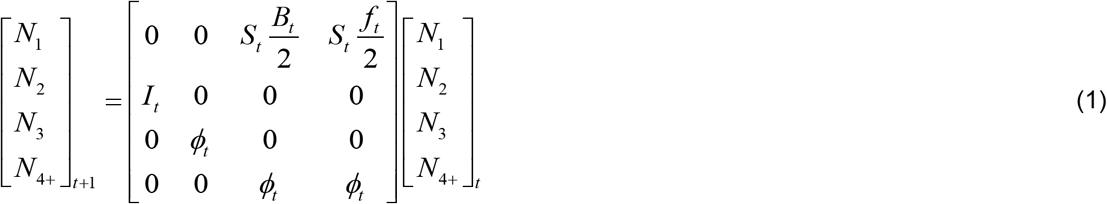

Here, birds survive the juvenile age class (0-1 years) with probability 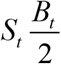 if the mother was age 3 years and 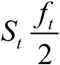 if the mother was age 4 or older, the immature age class (1-2 years) with probability *I*_*t*_ and one pre-breeding year (2-3 years) as well as two breeding age classes (3-4 years and all birds 4 years or older) with a survival rate equivalent to that of adults (*ϕ*_*t*_).

We ran the five PVA simulations for 1000 iterations. Within each iteration and across all five PVA simulations, temporal residuals associated with environmental stochasticity were held constant (Table 1) by using the same draws from the beta distribution for survival and gamma distribution for fecundity. We initiated each iteration with the number of breeding pairs (i.e. females) observed for kittiwakes breeding on Skomer Island in 1989 (N=2302) (Horswill et al. *in review*). Annual censuses of apparently occupied nests were conducted between 1989 and 2018 for kittiwakes breeding on Skomer Island. We ran each simulation for 30 years to match the duration of the count data available. The stable age distribution was derived from the dominant eigenvector of the Leslie matrix based on mean demographic rates (Caswell 2001). Following other assessments of seabird colonies that use a Leslie matrix modelling approach (Miller et al. 2019), we applied a maximum colony size to prevent exponential population growth (*N*_max_ = 2750). This threshold sets a carrying capacity at 448 breeding pairs above the starting colony size. All models included demographic stochasticity using binomial and Poisson distributions for survival and fecundity events, respectively, and all analysis was constructed and run using Program R (v. 4.0.2) (R Core Team 2020).

To examine predictive accuracy across the five PVA simulations (Table 1), we compared the simulated trajectories to the observed count data for kittiwakes breeding on Skomer Island. To evaluate how observed temporal trends in vital rates may influence the simulated impact associated with a hypothetical threat, we compared the output of the models with and without this demographic structure, as well as the additional 5% mortality, i.e., Model 2 vs. Model 3, and Model 4 vs. Model 5 (Table 1).

## Results

### Prevalence of temporal trends in vital rates

We identified 72 colonies of kittiwakes breeding in the UK with more than eight years of fecundity data between 1985 and 2019 (mean = 21 years, SD = 9 years). Of these, 79% (n=57) demonstrated a significant temporal trend in their rates of fecundity (Fig. 1, p-value ≤0.05). This included 46 colonies with a consistent decrease in fecundity, of these 70% (n=32) had a steeper slope coefficient than that used in the PVA simulations (national range of negative slope coefficients: -0.57, -1.83 ×10^−3^; slope coefficient for fecundity at Skomer: -0.02). There were also 11 colonies where fecundity consistently increased over time (range of positive slope coefficients: 6.79 ^x10-3^, 0.16), and 15 colonies with no trend (i.e., stable fecundity, Fig. 1). At the national level, the fecundity of kittiwakes demonstrated a significant negative temporal trend (slope estimate: 0.01, SE = 2.89×10^−4^, z = -39.27, p<0.001).

### How do temporal trends in vital rates influence predicted population dynamics?

We ran five PVA simulations (Table 1) to examine how incorporating observed temporal trends in vital rates may influence predictive accuracy, as well as the projected impact associated with a hypothetical threat. The population trajectory predicted by the deterministic model (Model 1, Table1) declined slightly but not at the rate observed in the colony (Fig. 2A). Likewise, the mean population trajectory predicted by the stochastic model (Model 2) overestimated colony size, such that the lower limit of the 95% confidence interval was above the observed trajectory (Fig. 2A). By contrast, the population trajectories simulated by the model including the observed decline in rates of fecundity (Model 4) were much closer to the observed count data (Fig. 2B). As expected, including an additional 5% annual mortality across all age classes increased the rate of population decline (Fig 2). Combining this hypothetical threat with the observed trend in fecundity also meant that local extinction was highly likely to occur within the 30 year timeframe of the simulation (Fig. 2B).

**Fig. 2.**
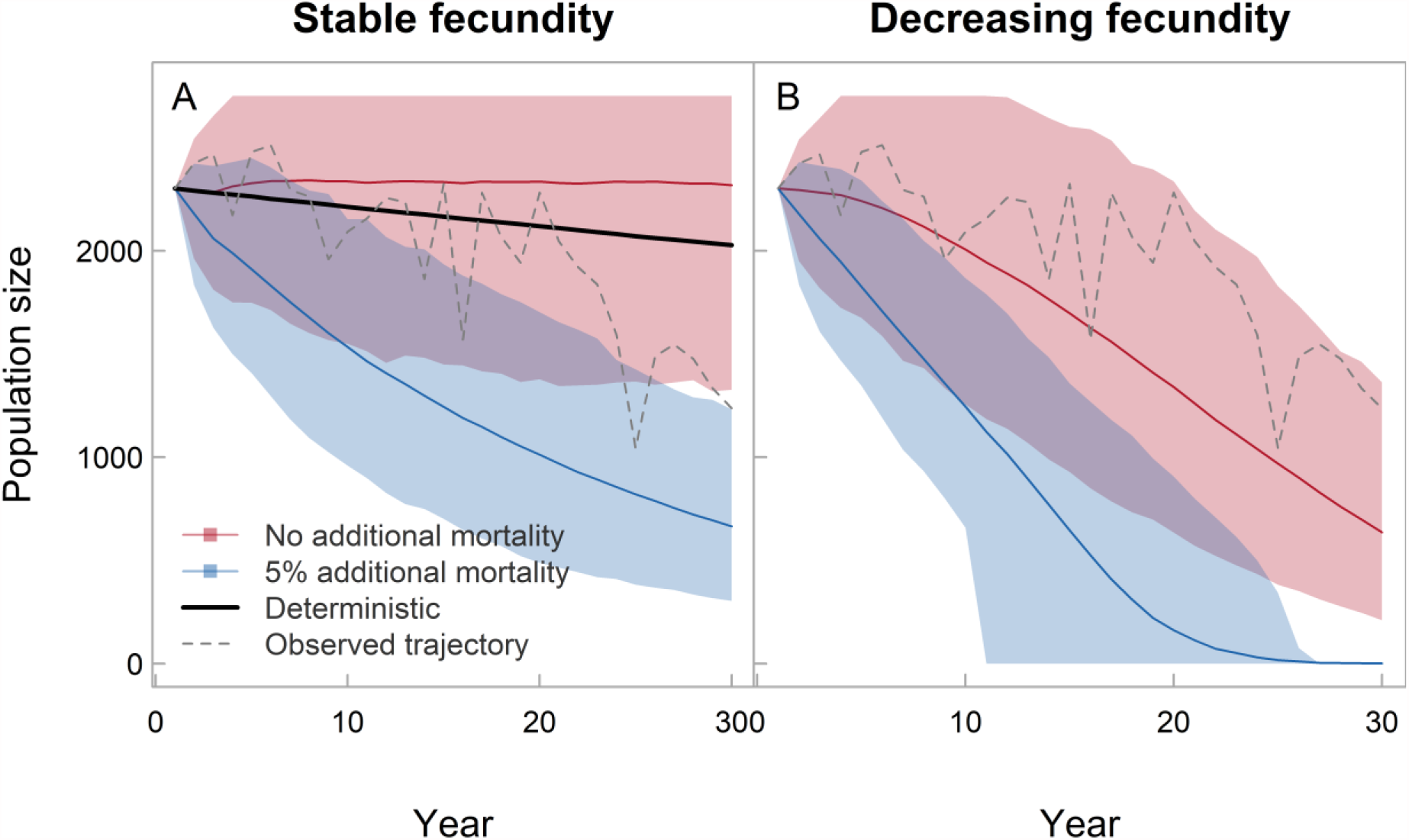
Projecting population dynamics without including observed temporal trends in vital rates will overestimate population size (A). Including temporal trends in fecundity improves the predictive accuracy of the model and ability to recreate the observed population trajectory (B). In both panels, the rate of population decline increases with the addition of a hypothetical threat (blue line and shading); this rate of decline is considerably faster when combined with the observed negative trend in fecundity (Panel B). Panel A shows Model 1 (Deterministic, black bold line), Model 2 (environmentally stochastic vital rates without additional mortality, red bold line shows mean trajectory and red shading shows 95% confidence interval) and Model 3 (environmentally stochastic vital rates with additional 5% mortality, blue bold line shows mean trajectory and blue shading shows 95% confidence interval). Panel B shows Model 4 (environmentally stochastic vital rates and trended fecundity without additional mortality, red bold line shows mean trajectory and red shading shows 95% confidence interval) and Model 5 (environmentally stochastic vital rates, trended fecundity and additional 5% mortality, blue bold line shows mean trajectory and blue shading shows 95% confidence interval). Observed population trajectory for kittiwakes at Skomer shown as grey dashed line in both panels.

## Discussion

In an era of global climate change and habitat degradation (Maxwell et al. 2016), many populations of wild animals are experiencing temporal trends in their vital rates, such as survival and fecundity (e.g. Gaillard et al. 2013; Sydeman et al. 2021, Fig. 1). However, this demographic structure is often overlooked in impact assessments that evaluate potential risks to threatened populations (Chaudhary & Oli 2020; Searle et al. 2020). In this study, we show how the predictive accuracy of population viability analysis (PVA) is greatly improved by incorporating observed trends in vital rates and demonstrate that omitting this structure can result in threats being underestimated.

To recreate the observed population trajectory for kittiwakes breeding on Skomer Island, it was necessary to include a negative trend in rates of fecundity. The deterministic model failed to replicate the observed rate of decline, as did the stochastic model with a stable mean rate of fecundity. This result emphasises that an essential prerequisite step to using PVA for impact assessment should be testing the predictive accuracy of models by simulating dynamics for a period with available count data. For populations where vital rate data are available, a mismatch between the simulated and observed trajectories will highlight that the distribution of vital rates (mean or variance) has changed over the study period, and that this demographic structure needs to be incorporated to support reliable assessments of impact.

In line with observations for kittiwakes breeding on Skomer Island, we applied the temporal trend to rates of fecundity. Previous studies show that in kittiwakes, like other long-lived species, fecundity has low demographic impact on population dynamics, i.e. low elasticity (Horswill et al. 2021). However, environmental stochasticity can increase the demographic impact of vital rates with low elasticity (Gaillard et al. 1998), and we show that a directional trajectory enables fecundity to play a predominant role in population dynamics. It would be understandable to assume that accurately quantifying vital rates with low demographic impact is of low priority when parameterising PVAs. However, our analysis demonstrates that this assumption could lead to erroneous impact assessments and misinformed management decisions.

We show that temporal trends in rates of fecundity are common and widespread for kittiwakes breeding in the UK (Fig. 1). A recent meta-analysis demonstrates that this trend is reflected in fish-eating, surface-foraging species of seabird, like kittiwakes, across the northern hemisphere (Sydeman et al. 2021). In our case study colony, fecundity is declining at a shallower rate than observed in many of the other UK colonies of kittiwakes. For the colonies that demonstrate a greater rate of change in fecundity, omitting temporal trends from impact assessments is likely to have a greater effect than demonstrated here. However, not all populations will have long-term monitoring data available. When conducting impact assessments on data-limited populations, we recommend an appropriate monitoring study be undertaken to quantify the local population trajectory and stability of demographic rates. If this is not possible, an alternative approach is to reconstruct mean vital rates using predictive methods based on life-history trade-offs (Horswill et al. 2019, 2021) and examine the potential consequences for population dynamics if vital rates change over time according to a realistic range of slope coefficients, as seen in other local populations of the same species. Interpreting the results of this latter approach may be complicated by large confidence intervals.

We focus on the implications of overlooking temporal trends in vital rates when parameterising impact assessment PVA models. Reviews of existing PVA literature identify a range of other considerations for applying this method to impact assessments and conservation decision making (e.g. Beissinger & Westphal 1998; Reed et al. 2002; Norris 2004; Pe’er et al. 2013; Chaudhary & Oli 2020; Searle et al. 2020). Other extrinsic factors that are often overlooked in impact assessments include density-dependent regulation. Depensatory and compensatory regulation are reported in colonies of kittiwakes (Horswill et al. 2017), however including density-dependent regulation in impact assessment models for seabirds remains debated (Green et al. 2016; Horswill et al. 2017; O’Brien et al. 2017). Consequently, there is a pressing need for more research determining the prevalence of density-dependence feedbacks across these species, how best to incorporate these structures into impact assessment models and the sensitivity of results to their inclusion.

In this paper, we demonstrate that temporal trends in rates of fecundity reflecting climate change and habitat degradation are relatively widespread for populations of kittiwakes breeding in the UK. We also show that omitting this structure from PVA models used to assess potential impacts to protected populations can dramatically reduce predictive accuracy and underestimate projected risk. We advocate that a prerequisite step to conducting impact assessments with PVA models involves testing predictive accuracy by simulating population dynamics for a period with available count data. If simulated dynamics do not match the observed trajectory, then incorporating a change in the distributions of vital rates will be critical for obtaining reliable results. More broadly, this study highlights the importance of incorporating long term climate and environmental change scenarios into conservation decision making.

## Acknowledgments

This work was funded by Research England to CH, and by the UK Joint Nature Conservation Committee (DEFRA)’s Seabird Monitoring Programme to MJW. We thank the Wildlife Trust of South and West Wales for logistical support and permission to work on Skomer Island, and the many fieldworkers who collected monitoring data. We also thank the Seabird Monitoring Program partners that provide and support the long-term monitoring of seabirds at a national scale.

